# Modelling longitudinal binary outcomes with outcome-dependent observation times: an application to a malaria cohort study

**DOI:** 10.1101/252411

**Authors:** Levicatus Mugenyi, Sereina A. Herzog, Niel Hens, Steven Abrams

## Abstract

Inspite of the global reduction of 21% in malaria incidence between 2010 and 2015, the disease still threatens many lives of children and pregnant mothers in African countries. A correct assessment and evaluation of the impact of malaria control strategies still remains quintessential in order to eliminate the disease and its burden. Malaria follow-up studies typically involve routine visits at pre-scheduled time points and/or clinical visits whenever individuals experience malaria-like symptoms. In the latter case, infection triggers outcome assessment, thereby leading to outcome-dependent sampling (ODS). Ordinary methods used to analyse such longitudinal data ignore ODS and potentially lead to biased estimates of malaria-specific transmission parameters, hence, inducing an incorrect assessment and evaluation of malaria control strategies. In this paper, we propose novel methodology to handle ODS using a joint model for the longitudinal binary outcome measured at routine visits and the clinical event times. The methodology is applied to malaria parasitaemia data from a cohort of *n* = 988 Ugandan children aged 0.5–10 years from 3 regions (Walukuba – 300 children, Kihihi – 355 children and Nagongera – 333 children) with varying transmission intensities (entomological inoculation rate equal to 2.8, 32 and 310 infectious bites per unit year, respectively) collected between 2011–2014. The results indicate that malaria parasite prevalence and force of infection (FOI) increase with age in the region of high malaria intensities with FOI highest in age group 5–10 years. For the region of medium intensity, the prevalence slightly increases with age and the FOI for the routine process is highest in age group 5–10 years yet for the clinically observed infections, the FOI gradually decreases with increasing age. For the region with low intensity, both the prevalence and FOI peak at the age of one year after which the former remains constant with age yet the latter suddenly decreases with age for the clinically observed infections. In all study sites, both the prevalence and FOI are highest among previously asymptomatic children and lowest among their symptomatic counterparts. Using a simulation study inspired by the malaria data at hand, the proposed methodology shows to have the smallest bias, especially when consecutive positive malaria parasitaemia presence results within a time period of 35 days were considered to be due to the same infection.

## Introduction

Malaria is potentially life-threatening and infections are caused by Plasmodium parasites that are transmitted through bites of infected female mosquitoes. In spite of the fact that malaria is a preventable and curable disease for which increased efforts worldwide dramatically reduced malaria incidence (i.e., a reduction of 21% between 2010 and 2015 as reported by WHO [1]), African countries still carry a disproportionately high share of the overall malaria burden. In order to reduce the malaria burden in African countries such as Uganda, a correct assessment and evaluation of the impact of control strategies is quintessential. Measures of malaria transmission intensity such as the entomological inoculation rate (EIR), the parasite prevalence and the malaria force of infection (FOI) have been used frequently to quantify the impact of various interventions [2,3]. In general, malaria transmission has been reported to be highly inefficient, meaning that the ratio of EIR to FOI is relatively high. As is the case for other infections, individual-and household-specific heterogeneity in malaria acquisition is hardly ever accounted for in the estimation of the aforementioned epidemiological parameters, albeit that it is well-recognised that variability in environmental and host-related factors, among other sources, has an important effect thereon [4].

Often in clinical trials with follow-up to study (infectious) disease dynamics, study participants are asked to come to the clinic and get examined for malaria infection during scheduled (routine) visits. On top of that, unscheduled (clinical) visits can occur when participants develop symptoms for the disease under consideration, or when they experience symptoms similar to those typically observed for the infection at hand. If infection triggers outcome assessment in between prescheduled follow-up visits, the outcome and observation-time processes are said to be dependent, which in literature is often referred to as outcome-dependent sampling (ODS) [5]. Conventional longitudinal methods to analyse repeated measurements for subjects over time assume independence of both processes. Hence, such unscheduled visits, and the ODS they induce, could lead to biased estimation of the epidemiological quantities of interest when not appropriately accounted for in the statistical analysis.

Different models have been proposed to address ODS in different experimental settings. For example, Ryu *et al*. [6] considered studies where the measurement time points are unequally spaced and having a follow-up measurement at any time depends on the history of past visits and outcomes of that individual. These authors discussed limitations of previously proposed models and methods for longitudinal data, such as generalised linear mixed models and generalised estimating equations (GEE), which do not address the association between the outcome and observation time process. Furthermore, these authors proposed a joint model using latent random variables in which the observed follow-up times are described together with the longitudinal response data [6]. More recently, Tan [5] considered a joint model with a semi-parametric regression model for the longitudinal outcomes and a recurrent event model for the observation times. Rizopoulos *et al*. [7] stated that an attractive paradigm for the joint modelling of longitudinal and time-to-event processes is the shared parameter framework [11] in which a set of random-effects is assumed to induce the interdependence of the two processes.

Although several authors developed methods to accommodate ODS in various settings, we propose new methodology to cope with both routine and clinical data on malaria infections from a cohort study in Uganda. More specifically, this paper focuses on the estimation of the malaria parasite prevalence in three regions of Uganda, accounting for observed and unobserved heterogeneity as done previously, while dealing with ODS at the same time. The paper is organised as follows. Our motivating example is introduced and briefly discussed in Section 2. In Section 3, we present the general methodology to estimate malaria FOI from parasitaemia data. In Section 4, we briefly highlight the impact of ignoring ODS after which our proposed joint model is fitted to the available routine and clinical data on parasite presence in Ugandan children in Section 5. Finally, these results are discussed in Section 6 together with strengths and limitations of the proposed methodology.

## Motivating example

In this paper, we consider longitudinal cohort data from children aged 0.5 to 10 years in three regions in Uganda; Nagongera sub-county, Tororo district; Kihihi sub-county, Kanungu district; and Walukuba sub-county, Jinja district. The data were collected as part of the Program for Resistance, Immunology, Surveillance and Modelling of malaria (PRISM) study [3]. The aforementioned study regions are characterized by distinct transmission intensities, with the highest intensity reported in Nagongera, followed by Kihihi and with Walukuba having the smallest intensity [3, 4]. The study participants were recruited from 300 randomly selected households (100 per region) located within the catchment areas. In total, *n* = 988 children were followed over time with 300 children in Walukuba, 355 in Kihihi and 333 children in Nagongera. Individuals were routinely tested for the presence of Plasmodium parasites using microscopy every three months from August 2011 to August 2014 (3 years). Furthermore, tests were also conducted at unscheduled clinical visits. More detailed information regarding the study design can be found in Kamya *et al*. [3].

Throughout this paper, the outcome process refers to the occurrence of the longitudinal binary outcome (parasite presence), and the observation-time process relates to the timing of scheduled, i.e. routine, and unscheduled, i.e. clinical visits over the entire follow-up period of the study.

## Materials and methods

### 3.1 Malaria dynamics – A simplified transmission model

For the purpose of this paper, we consider a simplified version of a realistic transmission model to describe malaria infection dynamics. More specifically, following Mugenyi *et al*. [4], a so-called Susceptible (S) - Infected (I) - Susceptible (S), or short SIS, compartmental model dividing the population into two mutually exclusive compartments, i.e., the susceptible (S) and infected (I) class, will be used to describe malaria dynamics within the human host. We refer to the discussion of [4] for a motivation of the choice of the SIS model and would like to note that the methodology outlined here is more generally applicable in case of other disease dynamics. The schematic diagram depicting the flows between the different states is graphically displayed in Figure 1.

**Figure 1:**
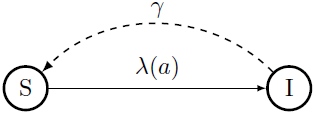
A schematic diagram of the SIS compartmental model illustrating the simplified dynamics for malaria transmission: Individuals are born into the susceptible class S and move to the infected state I at age-specific rate λ(*a*), after which they become susceptible again at rate *γ*.

Herein, the force of infection λ(*a*) represents the instantaneous rate at which individuals flow from the susceptible compartment S to the infected state I at age *a*, i.e., the age-specific rate at which individuals are infected with malaria parasites through effective mosquito bites. Furthermore, *γ* represents a time- and age-invariant clearance rate at which individuals regain susceptibility after clearing malaria parasites from their blood. Let *s*(*a*) denote the proportion of susceptible individuals in the population and *i*(*a*) the proportion of infected individuals of age *a*, i.e., the (point) parasite prevalence, then the following set of ordinary differential equations (ODEs) describes transitions in the compartmental SIS model:

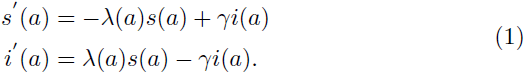

Hence, one can easily derive the following expression for the age-dependent force of infection in terms of the point prevalence *i*(*a*):

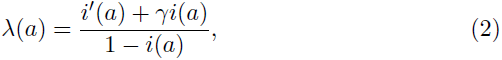

using *i*(*a*) + *s*(*a*) = 1. Hence, constructing a model for the point prevalence *i*(*a*) will imply a specific functional form for the underlying force of infection λ(*a*) depending on the clearance rate γ.

### 3.2 Parasite prevalence and routine visits

Consider the binary random variable *Y*_*ij*_ representing an indicator for the presence of malaria parasites for individual *i* at (routine) visit *j*. Consequently, for scheduled routine visits, 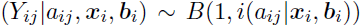, where *a*_*ij*_ represents the age of individual *i* at visit *j*, *x*_*i*_ represents a (*p* × 1)-vector of covariate information for individual *i* = 1, …, *n*, and *b*_*i*_ a (*q* × 1)-vector of individual-specific random effects. In order to model the parasite prevalence, we formulate a generalized linear mixed model with cloglog-link as follows:

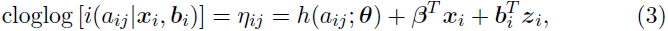

where *β* is a column vector of unknown regression parameters and *z*_*i*_ is an individual-specific (*q* × 1) design vector for ***b***_*i*_ which is a column vector of individual-specific normally distributed random effects, i.e., ***b*_*i*_** ~ *N*(***μ*, *D***) thereby addressing the association among repeated measurements over time within the same individual. Here, the variance-covariance matrix *D* is assumed to have zero elements, except for the the variances on the main diagonal. Moreover, *h*(*a*_*ij*_; ***θ***) is a known function describing the age-effect with parameter vector ***θ***. Note that the calendar time effect can be introduced in the linear predictor by means of the shifted birth year of the *i*th individual, implying the prevalence, and equivalently the FOI, to depend on both age and calendar time [4]. In Table 1, we present some common parametric distributions and their implied functional forms for *h*(*a*_*ij*_; ***θ***) based on model (3) and the corresponding baseline infection risk 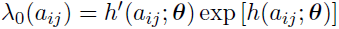(derived under the assumption of no parasite clearance).

In the absence of unscheduled clinical visits (*n*_*i*_ = *n*_*i(r*)_, i.e., the number of routine visits for individual *i*), or under the assumption of independence between the observation time process and the outcome process, we can simply estimate model parameters using maximum likelihood techniques, thereby maximizing a marginal likelihood function with the following individual likelihood contributions:

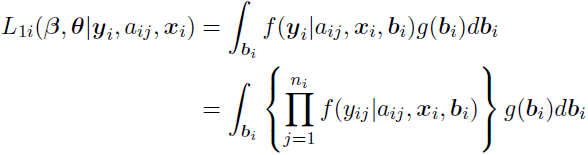

with

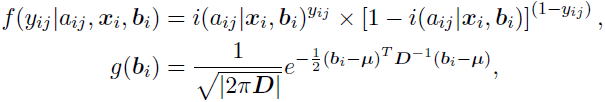

where *y*_*ij*_ is the observed binary outcome for individual *i* at routine visit *j* = 1, …, *n*_*i*_, and 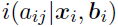 is the conditional parasite prevalence. Numerical integration techniques are employed to perform integration over the random effects distribution *g*(***b***_*i*_). In the following subsection, we specifically focus on clinical visits and how to address ODS.

**Table 1:** Distributional assumptions regarding the underlying age-specific malaria force of infection.

| Distribution | $\theta$ | $h(a_{ij}; \theta)$ | $\lambda_0(a_{ij})$ |
| --- | --- | --- | --- |
| Exponential | $\theta_1 > 0$ | $\log(\theta_1 a_{ij})$ | $\theta_1$ |
| Weibull | $\theta_1, \theta_2 > 0$ | $\log(\theta_1 a_{ij}^{\theta_2})$ | $\theta_1 \theta_2 a_{ij}^{\theta_2 - 1}$ |
| Gompertz | $\theta_1 > 0, -\infty < \theta_2 < +\infty$ | $\log \left[ \frac{\theta_1}{\theta_2} (e^{\theta_2 a_{ij}} - 1) \right]$ | $\theta_1 e^{\theta_2 a_{ij}}$ |
| Log-logistic | $\theta_1, \theta_2 > 0$ | $\log \left\{ \log \left[ 1 + (\theta_1 a_{ij})^{\theta_2} \right] \right\}$ | $\frac{\theta_1 \theta_2 (\theta_1 a_{ij})^{\theta_2 - 1}}{1 + (\theta_1 a_{ij})^{\theta_2}}$ |
| Fractional polynomial | $\theta_2 < 0$ | $\theta_2 a_{ij}^{-1}$ | $-\theta_2 a_{ij}^{-2} e^{\theta_2 a_{ij}^{-1}}$ |

## 3.3 Outcome-dependent sampling and clinical visits

As mentioned before, clinical visits due to symptomatic malaria infections, or malaria-like events giving rise to symptoms similar to those observed for malaria, can not be treated in the same way as described in Section 3.2. Let *t*_*ij*_ represents the time-at-risk for an individual *i* for which the *j*th visit is clinical, and *c*_*ij*_ an indicator having value one for an unscheduled clinical visit and 0 for routine data. For the purpose of illustration, we assume that *t*_*ij*_ is known, albeit that this is not the case in practice, and statistical ways to deal with this are outlined below. The probability density function for the random variable *T*_*ij*_, suppressing dependence on covariates *x*_*i*_ and *c*_*ij*_ = 1 for simplicity, is given by:

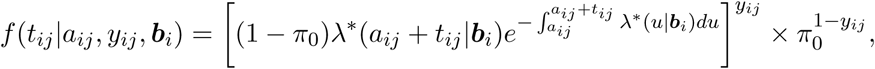

where 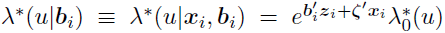 is the conditional time-varying malaria force of infection under the proportional hazards assumption (with ζ a vector of model parameters) and π_0_ denotes the probability of a malaria-like clinical visit for which no malaria parasites are present in the blood. For the purpose of this paper, we will not model the dependence of the probability of having a malaria-like event 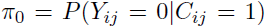on the observed covariate information *a*_*ij*_ and *x*_*i*_. Different distributional assumptions can be made regarding the time-at-risk distribution, such as, e.g., exponential, Weibull, Gompertz, among others, which also relates to the selected functional form for *h*(*a*_*ij*_; ***θ***) in the outcome process model (see Section 3.2 and Table 1). In order to align the models for both processes, the baseline infection risk 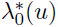 for the observation time process can be of the same type as λ_0_(*u*), albeit that distributional parameters, say *ϑ*, are allowed to be different. Note that more flexible parametric shapes for *h*(*a*_*ij*_; ***θ***), such as, e.g., using fractional polynomials, could result in nonstandard non-negative distributions for the malaria infection times, albeit that unconstrained optimisation could lead to negative FOI estimates. In the statistical analyses, we include parametric fractional polynomials as an alternative to the standard event time distributions.

For outcomes 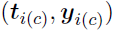that are derived from the clinical visits *j* = 1, …, *n*_*i*(*c*)_, where 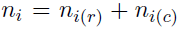 having *n*_*i*(*r*)_ and *n*_*i*(*c*)_ the number of routine and clinical visits for individual *i*, respectively, the likelihood function has contributions:

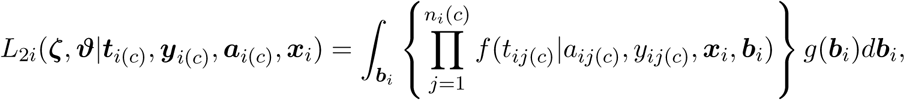

Where *t*_*i*(*c*)_ and *a*_*i*(*c*)_ are the vectors of time-at-risk and age values at which the individual becomes at risk for the *j*th clinical event, respectively. Finally, the likelihood for the joint model including both information on routine and clinical visits is obtained by combining likelihood contributions as described before:

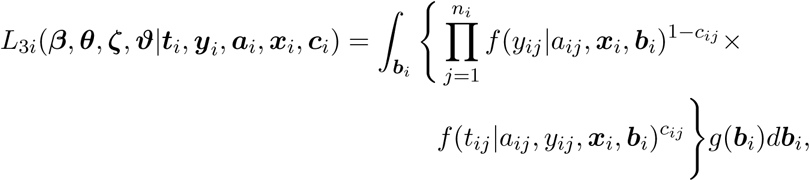

At least under the assumption that each malaria event contributes solely to one of the two components (i.e., routine or clinical process) in the likelihood. As mentioned previously, the time-at-risk for a specific clinical event (i.e., a symptomatic malaria infection) is not precisely known. More specifically, malaria infection times are interval-censored which needs to be taken into account in the statistical analyses through the modification of the likelihood function. For more details on how the interval-censoring has been treated in the analyses, the reader is referred to Appendix B.1.

## Simulation study

In order to study the impact of ignoring ODS, we set up a simulation study which is inspired by the PRISM data under consideration. More specifically, we generate *M* = 1000 datasets including *n*_*m*_ ≡ *n* = 1000 individuals per simulated dataset (*m* = 1, …, *M*). Furthermore, we consider a simulation setting in which exponential infection times occur during a follow-up period of 1800 days (≈ 5 years) and with an average duration until acquiring a new infection of about 365 days (1 year: 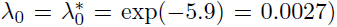Parasite clearance times are exponentially distributed with a mean duration of infectiousness equal to 50 days (*γ* = 0:02). Based on the generated infection histories for the individuals, routine and clinical visits are obtained. More specifically, routine visits are scheduled every 90 days and parasite presence is recorded based on the current status at the time of data collection. Varying probabilities for having a symptomatic malaria episode are considered in the simulation whereby symptomatic observations at unscheduled time points were considered as clinical visits (i.e., *P* = 20%; 40%; 60%; 80%; 100%). Hence, asymptomatic malaria cases were only included when detected during the routine process. No malaria-like events were generated such that all clinical visits are due to symptomatic malaria infections (i.e., *π*_0_ = 0). Individual-specific random intercepts 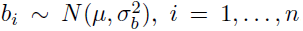with 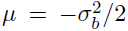implying a unit mean for the lognormal random terms *e^b_i_^*, are introduced to induce correlation between repeated measurements for the same subject 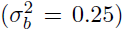 If a single infection is contributing to both the routine and clinical process (i.e., consecutive observations *C*^+^ and *R*^+^, or vica versa), hence leading to two dependent observations, we drop the second one in Scenario 4. However, without additional information, we cannot determine whether individuals already recovered and got re-infected in between such visits, thereby potentially underestimating the FOI. We performed a sensitivity analysis given the simulation scenario at hand in order to deduce the time period in which consecutive positive routine and clinical observations can be considered to be the result of a single malaria infection. From this exercise, a period of 35 days is assumed to be optimal (see Appendix A, Figure A.1 for more details thereon). This observation is supported by the literature where 100% recovery rate was reported on day 28 following anti-malaria treatment [14, 15].

**Table 2:**
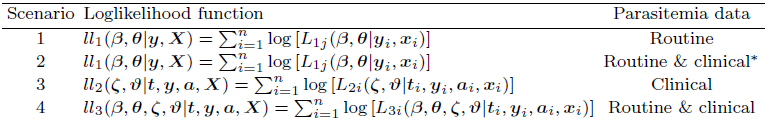
Overview of the different scenarios, corresponding loglikelihood functions to be maximised (see Section 3) and parasitaemia data that is included in the analyses. ^*^ Scenario does not take ODS into account.

## 4.1 Simulation results

Hereunder, we present results from fitting the four scenarios based on the three different likelihoods in Section 3 to the simulated data. All models converged for all simulation runs. In Table 3, we show the simulation results for the four different scenarios described in Table 2 with varying percentages of symptomatic malaria infections. Scenario 2 including both routine and clinical data without accounting for ODS performs worse compared to Scenario 1 in which only routine data is used. Hence, ignoring ODS leads to biased estimates of both the baseline hazard as well as population-averaged hazard functions. Note that Scenario 1 is not influenced by the percentage of symptomatic infections, simply since these clinical infections are not accounted for therein. Our proposed model for the analysis of both clinical and routine parasitaemia data (Scenario 4) outperforms Scenarios 1 and 2 in terms of bias and precision (and consequently MSE) for the baseline hazard function and population-averaged hazard λ_*p*_, at least when *P* = 60% or higher, and leads in all cases to a reduction in bias. In Scenario 4, we add clinical information to the readily available routine data (i.e., larger sample size), resulting in a lower MSE, bias and empirical variance for the model parameters compared to Scenario 1. The loss of perfomance in Scenario 4 compared to Scenario 3 as *P* > 60% can be explained by the nature of the data since noise is added by combining time-to-event data (which is analysed separately in Scenario 3) with interval-censored data.

**Table 3:** Simulaton results for the different models showing mean estimates for the marginal or population-averaged FOI (λ_*p*_), variance of the random intercepts 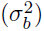, and the corresponding mean squared error (MSE), bias and empirical variance. *P* represents the percentage of symptomatic infections. ^†^: all data except for positive routine observations following a positive clinical visit, or positive clinical observations following a positive routine visit within a 35 day period. *N* represents the total number of observations over all individuals averaged over the *M* datasets.

|  | Scenario 1<br>routine data | Scenario 2<br>all data | Scenario 3<br>clinical data | Scenario 4<br>all data<sup>†</sup> |
| --- | --- | --- | --- | --- |
| $P = 20\%$ | $N = 20,000$ | $N = 21,832$ | $N = 1,832$ | $N = 21,781$ |
| **Population-averaged hazard $\lambda_p$** |  |  |  |  |
| Mean estimate $\bar{\lambda}_p$ | 0.0034 | 0.0047 | 0.0020 | 0.0028 |
| Mean estimate $\bar{\sigma}_b^2$ | 0.3806 | 0.4058 | 0.4727 | 0.5228 |
| MSE($\lambda$)$\times 10^5$ | 0.0078 | 0.2460 | 0.1375 | 0.0111 |
| Bias($\lambda$)$\times 10^5$ | 23.8252 | 155.6670 | 116.2956 | 9.0705 |
| Var($\lambda$)$\times 10^5$ | 0.0021 | 0.0037 | 0.0023 | 0.0102 |
| $P = 40\%$ | $N = 20,000$ | $N = 22,678$ | $N = 2,678$ | $N = 22,576$ |
| **Population-averaged hazard $\lambda_p$** |  |  |  |  |
| Mean estimate $\bar{\lambda}_p$ | 0.0034 | 0.0060 | 0.0026 | 0.0028 |
| Mean estimate $\bar{\sigma}_b^2$ | 0.3806 | 0.4046 | 0.3646 | 0.4402 |
| MSE($\lambda$)$\times 10^5$ | 0.0078 | 0.8106 | 0.0270 | 0.0093 |
| Bias($\lambda$)$\times 10^5$ | 23.8252 | 283.8542 | 50.4948 | 7.0972 |
| Var($\lambda$)$\times 10^5$ | 0.0021 | 0.0048 | 0.0016 | 0.0088 |
| $P = 60\%$ | $N = 20,000$ | $N = 23,520$ | $N = 3,520$ | $N = 23,370$ |
| **Population-averaged hazard $\lambda_p$** |  |  |  |  |
| Mean estimate $\bar{\lambda}_p$ | 0.0034 | 0.0072 | 0.0029 | 0.0028 |
| Mean estimate $\bar{\sigma}_b^2$ | 0.3806 | 0.3908 | 0.3076 | 0.3861 |
| MSE($\lambda$)$\times 10^5$ | 0.0078 | 1.6500 | 0.0063 | 0.0081 |
| Bias($\lambda$)$\times 10^5$ | 23.8252 | 405.4125 | 22.5147 | 5.4008 |
| Var($\lambda$)$\times 10^5$ | 0.0021 | 0.0064 | 0.0012 | 0.0078 |
| $P = 80\%$ | $N = 20,000$ | $N = 24,368$ | $N = 4,368$ | $N = 24,169$ |
| **Population-averaged hazard $\lambda_p$** |  |  |  |  |
| Mean estimate $\bar{\lambda}_p$ | 0.0034 | 0.0084 | 0.0030 | 0.0028 |
| Mean estimate $\bar{\sigma}_b^2$ | 0.3806 | 0.3780 | 0.2713 | 0.3496 |
| MSE($\lambda$)$\times 10^5$ | 0.0078 | 2.7799 | 0.0017 | 0.0072 |
| Bias($\lambda$)$\times 10^5$ | 23.8252 | 526.4963 | 8.4274 | 4.2648 |
| Var($\lambda$)$\times 10^5$ | 0.0021 | 0.0064 | 0.0012 | 0.0064 |
| $P = 100\%$ | $N = 20,000$ | $N = 25,218$ | $N = 5,218$ | $N = 24,971$ |
| **Population-averaged hazard $\lambda_p$** |  |  |  |  |
| Mean estimate $\bar{\lambda}_p$ | 0.0034 | 0.0072 | 0.0029 | 0.0028 |
| Mean estimate $\bar{\sigma}_b^2$ | 0.3831 | 0.3659 | 0.2445 | 0.3265 |
| MSE($\lambda$)$\times 10^5$ | 0.0083 | 4.1944 | 0.0007 | 0.0065 |
| Bias($\lambda$)$\times 10^5$ | 23.8252 | 646.9211 | 1.0401 | 3.5431 |
| Var($\lambda$)$\times 10^5$ | 0.0021 | 0.0064 | 0.0012 | 0.0064 |

### Data application

In this section, we apply the proposed joint model in Scenario 4 to the observed Ugandan malaria parasitaemia data presented in Section 2. The covariates considered in the model building process included study site, age, shifted birth year (i.e., shifted birth year = birth year - birth year of the oldest child), previous use of Artemether-Lumefantrine (AL) treatment, and the infectious status at the previous visit. The covariate ‘shifted birth year’ was generated to represent the calendar time (see also [4] for details concerning this modelling strategy). Let *S*_*i*_ represent the study site (1 = Walukuba, 2 = Kihihi, 3 = Nagongera), *a*_*ij*_ the child’s age in years, *l*_*ij*_ the shifted birth year, *P_ij_* the previous infection status and use of AL (1 = Negative & no AL, 2 = Negative + AL, 3 = Symptomatic, 4 = Asymptomatic) for individual *i* at visit *j*. Different parametric distributional assumptions regarding the infection times are explored (i.e., leading to various functional forms for *h*(*a*_*ij*_; ***θ***), and equivalently, for the underlying malaria force of infection) thereby allowing for different distributional parameters θ and ϑ for the outcome and infection time process, respectively. Since malaria transmission intensity differs between the three sites (see, e.g., [3,4]), site-stratified analyses were performed, and model comparison was done based on *AIC* and *BIC* in order to select the most appropriate functiontal form for *h*(*a*_*ij*_; ***θ***). Table B.1 in Appendix B provides the site-specific fit statistics for the different models.

In Table 4, we show the parameter and standard error estimates (between brackets) for the joint model under Scenario 4, thereby having Gompertz baseline hazard functions λ_0_(a) and 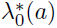 for the three study sites (see Table B.1 in Appendix B for more details on the *AIC*- and *BIC*-values for the candidate models). A signi cant effect of shifted year of birth has been observed for Kihihi and Nagongera in both processes, and not for the low transmission intensity site Walukuba. The infection status at previous visit was included only for the outcome process resulting in an overall significant effect at all sites (p-value <0.001). In total, 35%, 43% and 62% of the observed visits were classified as clinical visits in Walukuba, Kihihi and Nagongera, respectively. Of those observed clinical visits, 87%, 48% and 54% are malaria-like clinical visits implying that no evidence of malaria infection was found in children coming to the clinic due to malaria-like symptoms. The estimated values for π_0_ are equal to 89% (95% confidence interval (CI): 87% - 91%), 58% (95% CI: 56% - 60%) and 65% (95% CI: 63% - 67%) for Walukuba, Kihihi and Nagongera, respectively, which are quite in line with the observed empirical probabilities.

Figure 2 depicts the estimated marginal prevalence by age for children assumed to be born in the baseline year (2001) which were symptomatic (top row) or asymptomatic (bottow row) at the previous visit, and by study site (left to right: Nagongera, Kihihi and Walukuba). The curves are drawn for Scenario 2 (solid blue line) and Scenario 4 (dashed red line). In general, the parasite prevalence increases with increasing age in areas with high (Nagongera) and medium (Kihihi) transmission intensity, though the prevalence is fairly constant for Scenario 4 in the latter case. In Walukuba, the prevalence first increases to a plateau from 6 months up to 2 years after the prevalence remains constant. From the graphs, it is clear that small differences exist between the two scenarios in terms of the estimated marginal prevalence.

**Figure 2:**
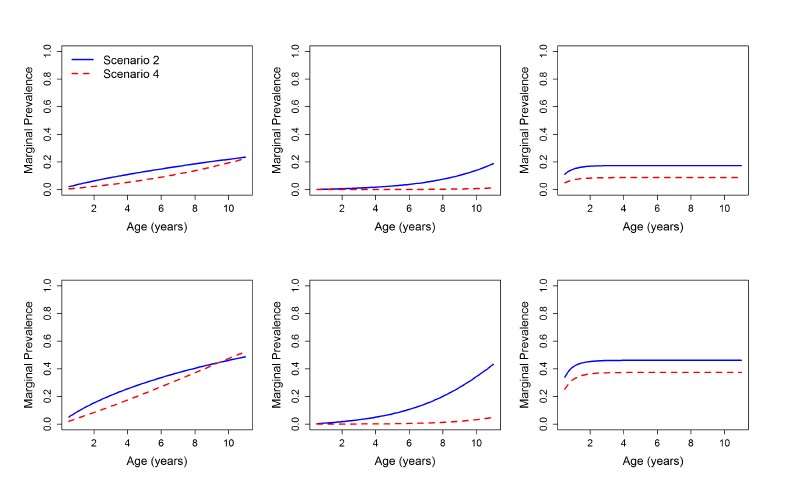
Estimated marginal prevalence for children assumed to be born in the baseline year (2001) by age, study site and symptomatic (top row) or asymptomatic (bottom row) at the previous visit. Left to right column: Nagongera, Kihihi and Walukuba.

In Figure 3, we show the estimated marginal FOI for the outcome (routine) process based on expression (2). We consider annual parasite clearance rates (γ) of 1.643, 0.584 and 0.986 years ^−1^ for children aged less than 1 year, 1-4 years and 5-10 years, respectively [19]. On top of that, the marginal FOI estimated from the time-to-event process is shown in the bottom row. The marginal FOI for the outcome process increases with increasing age at least for Nagongera and Kihihi, and it is highest among children in age group 5–10 years or those that were previously asymptomatic (gray bars) and least in their symptomatic counterparts (brown bars) in all study areas. For the time process, the marginal FOI in Nagongera is close to zero and constant with time at risk, at least for children aged 1 year when becoming at risk. For children at a higher age, the FOI tends to increase more steeply with increasing time at risk and age. However, the FOI for the time process is highest among children aged about one year in medium (Kihihi) and low (Walukuba) transmission intensities, after which it decreases gradually with increasing time at risk for children of all ages. More specifically, when children are older, the infection risk is smaller as compared to their younger counterparts given the specific time at risk.

**Figure 3:**
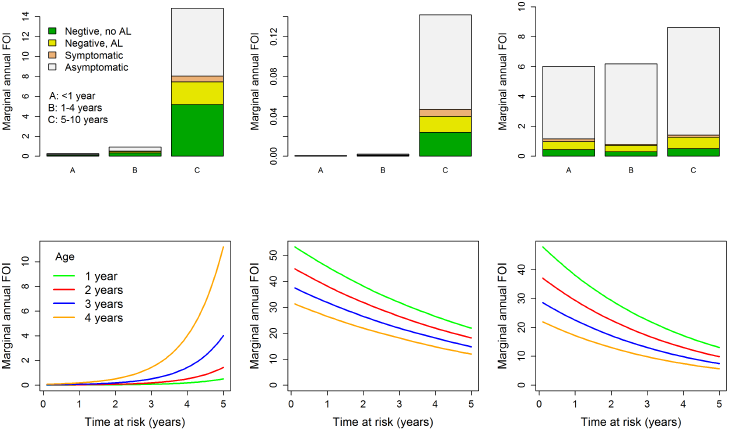
Estimated marginal FOI by time at risk and age when becoming at risk for the next malaria infection based on Scenario 4 and for children assumed to be born in the baseline year (2001) Top row: marginal FOI based on outcome process. Bottom row: marginal FOI based on time process. Left to right column: Nagongera, Kihihi and Walukuba.

**Table 4:** Application to PRISM data: results showing parameter and standard error (s.e.) estimates from the joint model (Scenario 4) assuming Gompertz-distributed infection times for Walukuba, Kihihi, and Nagongera.

| Effect |  | Parameter | Estimate (s.e.) | *t*-value | *p*-value |
| --- | --- | --- | --- | --- | --- |
| **Walukuba (Gompertz):** |  |  |  |  |  |
| Infection status at the previous visit (Ref = Negative & No AL treatment in past): | Negative + AL | $\beta_1$ | 0.12 (0.26) | 0.47 | 0.639 |
| | Symptomatic | $\beta_2$ | -0.66 (0.46) | -1.43 | 0.153 |
| | Asymptomatic | $\beta_3$ | 1.33 (0.26) | 5.13 | < 0.001 |
| Shifted year of birth | | $\beta_4$ | -0.09 (0.05) | -1.93 | 0.054 |
| Age | | $\theta_1$ | 0.16 (0.23) | 0.71 | 0.480 |
| | | $\theta_2$ | -1.54 (1.96) | -0.78 | 0.434 |
| Shifted year of birth<sup>*t*</sup> | | $\zeta$ | -0.23 (0.17) | -1.37 | 0.171 |
| Age<sup>*t*</sup> | | $\vartheta_1$ | 36.68 (63.44) | 0.58 | 0.564 |
| | | $\vartheta_2$ | -0.28 (0.14) | -2.01 | 0.046 |
| Probability of a malaria-like clinical visit | | $\pi_0$ | 0.89 (0.01) | 102.18 | < 0.001 |
| Variance for random intercepts for subjects | | $d_{11}$ | 0.25 (0.18) | 1.38 | 0.167 |
| Variance for random intercepts for households | | $d_{22}$ | 1.22 (0.37) | 3.32 | 0.001 |
| **Kihhi (Gompertz):** |  |  |  |  |  |
| Infection status at the previous visit (Ref = Negative & No AL treatment in past): | Negative + AL | $\beta_1$ | -0.30 (0.06) | -5.29 | < 0.001 |
| | Symptomatic | $\beta_2$ | -1.08 (0.14) | -7.94 | < 0.001 |
| | Asymptomatic | $\beta_3$ | 0.65 (0.12) | 5.37 | < 0.001 |
| Shifted year of birth | | $\beta_4$ | 0.49 (0.04) | 13.03 | < 0.001 |
| Age | | $\theta_1$ | 4e-6 (2e-6) | 2.61 | 0.009 |
| | | $\theta_2$ | 0.56 (0.04) | 13.57 | < 0.001 |
| Shifted year of birth<sup>*t*</sup> | | $\zeta$ | -0.25 (0.04) | -6.94 | < 0.001 |
| Age<sup>*t*</sup> | | $\vartheta_1$ | 18.06 (6.98) | 2.59 | < 0.001 |
| | | $\vartheta_2$ | -0.26 (0.03) | -7.91 | < 0.001 |
| Probability of a malaria-like clinical visit | | $\pi_0$ | 0.58 (0.01) | 53.58 | < 0.001 |
| Variance for random intercepts for subjects | | $d_{11}$ | 0.27 (0.05) | 9.83 | < 0.001 |
| Variance for random intercepts for households | | $d_{22}$ | 4.28 (0.35) | 12.30 | < 0.001 |
| **Nagongera (Gompertz):** |  |  |  |  |  |
| Infection status at the previous visit (Ref = Negative & No AL treatment in past): | Negative + AL | $\beta_1$ | -0.46 (0.12) | -3.80 | < 0.001 |
| | Symptomatic | $\beta_2$ | -1.24 (0.13) | -9.35 | < 0.001 |
| | Asymptomatic | $\beta_3$ | 0.15 (0.13) | 1.22 | 0.222 |
| Shifted year of birth | | $\beta_4$ | 0.19 (0.08) | 2.44 | 0.015 |
| Age | | $\theta_1$ | 0.02 (0.02) | 1.23 | 0.219 |
| | | $\theta_2$ | 0.17 (0.11) | 1.58 | 0.115 |
| Shifted year of birth<sup>*t*</sup> | | $\zeta$ | 0.92 (0.05) | 18.31 | < 0.001 |
| Age<sup>*t*</sup> | | $\vartheta_1$ | 6e-4 (3e-4) | 1.88 | 0.061 |
| | | $\vartheta_2$ | 1.03 (0.05) | 19.64 | < 0.001 |
| Probability of a malaria-like clinical visit | | $\pi_0$ | 0.65 (0.01) | 83.81 | < 0.001 |
| Variance for random intercepts for subjects | | $d_{11}$ | 0.94 (0.15) | 6.42 | < 0.001 |
| Variance for random intercepts for households | | $d_{22}$ | 0.37 (0.15) | 2.43 | 0.016 |

<sup>*t*</sup> Time-to-event model effects

## Discussion

In this paper, we have proposed novel methodology to account for outcome-dependent sampling (ODS) when estimating malaria transmission parameters such as, for example, the parasite prevalence and the force of infection (FOI) in case of longitudinal cohort data with routine (scheduled) and clinical (unscheduled) visits. A simulation study, inspired by parasitaemia data from a cohort of Ugandan children who were tested for malaria parasites (parasitaemia) during such visits, was conducted in which different parametric functions were considered to model the age-specific malaria prevalence and FOI while accounting for both observed and unobserved heterogeneity. The results clearly indicate that ignoring ODS leads to biased estimates for the marginal force of infection, hence, leads to an incorrect assessment and evaluation of malaria control strategies. We demonstrate that the bias can be reduced by using a joint model in which both outcome (routine) and observation-time (clinical) components are present. In order to reduce the bias, we propose to treat malaria events within a period of 35 days after a first malaria infection as being part of the same infection. This is supported by the results presented by Maiga *et al*. [14] and Ndiaye *et al*. [15].

The results show that both the malaria parasite prevalence and the FOI increase with increasing age in an area of high (Nagongera) transmision intensity. The FOI is highest in children aged 5–10 years and it becomes higher as children grow older or are at risk for a longer time. For an area with medium (Kihihi) transmision intensity, whereas the parasite prevalence and the FOI for the outcome process increase with increasing age, the FOI for the clinically observed infections (time process) is highest among children aged 1 year and it gradually decreases with increasing age and time at risk. In Walukuba which is an area of low transmission intensity, however, the prevalence and FOI at least for the time process peak at the age of about one year, after which the former remains constant while the latter decreases with increasing age and exposure time at least when based on the time process. Further, both the prevalence and FOI are highest among the children with asymptomatic infections, and lower among the symptomatic ones or the previously treated children. These results are in line with those reported previously by Mugenyi *et al*. [4]. The high prevalence and FOI estimated among the older children particularly in area with high transmission is in agreement with the work by Doolan *et al*. [17]. These authors show that children older than 5 years act as reservoirs for malaria parasites or asymptomatic infections and are rarely treated, hence leading to an increased infection risk. On the other hand, the decrease in the clinically observed infections (time process), that is FOI, as age increases in both the medium and low transmission intensities can be attributed to acquired immunity due to past infections or increase in age as discussed by Doolan *et al*. [17]. In our statistical analyses, we also estimated the probability of a malaria-like event 0 which were quite in line with the empirical proportions in the three regions. However, 0 also encompasses potential di erences in reporting among the regions as individuals with symptoms will not always visit the clinic.

One way to avoid bias in estimating the epidemiological parameters of interest is the use of routine data only. This approach has been demonstrated in the past [4]. However, our methodology allows for a proper integration of all clinical data, including malaria-like events, in the data analysis, thereby enabling the study of potential varying effects for symptomatic (detected at clinical visits) and asymptomatic (derived from routine data) infections. From our statistical analyses of the PRISM data, the hypothesis of differential age-effects for symptomatic and asymptomatic infections is highly supported as models forcing the effects to be the same are clearly outperformed by their unrestricted and more flexible counterparts. Though the estimated parasite prevalence is in line with the observed data, more flexible parametric or semi-parametric baseline hazard functions could be considered in both processes which is an interesting avenue for further research. Furthermore, Mugenyi *et al*. [4] used a generalized linear mixed model to model the observed parasite prevalence after which the force of infection is derived using equation (2). One of the shortcomings in this paper is the simplification of no parasite clearance when deriving the baseline hazard function for the time process. This could lead to an underestimation of the respective FOI. We consider this as an interesting avenue for future research.

The proposed joint model can be extended to have a shared parameter *Ψ* to model the dependence between the outcome and observation time processes through individual- and process-specific random effects *b_i1_* and *b_i2_*, respectively (see, e.g, [11]). In that way, one can allow the process-specific random effects to act at di erent levels. However, applying this approach to the PRISM data forced us to exclude the household-specific random effect for convergence reasons. The models presented in this paper (*Ψ* = 1) outperformed the ones with different process-specific individual-level random effects in all regions, except for Nagongera, and the significance of covariates was not altered (not shown here).

## Acknowledgements

A lot of thanks to the study participants and the PRISM study team including Moses Kamya, Grant Dorsey and Sarah Staedke for their permission to use the data. We acknowledge the PRISM grant (U19AI089674) by the National Institutes of Health (NIH). We thank the Vlaamse Interuniversitaire Raad (VLIR) for the PhD financial support that enabled the first author to complete this work. NH gratefully acknowledges support from the University of Antwerp scientific chair in Evidence-Based Vaccinology, financed in 2009-2017 by a gift from Pfizer and in 2016 by a gift from GSK.

## Appendix

### Appendix A: Simulation study

**Table A.1.**
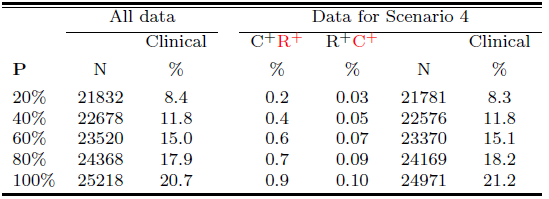
Average number of malaria episodes, by varying percentage of assumed symptomatic infections (P). The labels C^+^R^+^ and R^+^C^+^ represent positive results at two near-by visits (C = clinical and R = routine) with the second observation deleted.

**Figure A.1.**
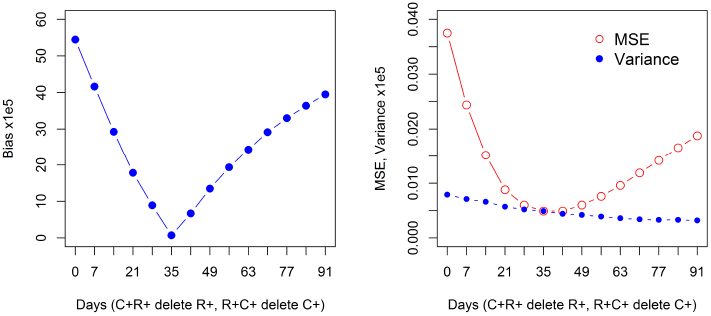
Sensitivity analysis for bias, MSE and variance obtained using Scenario 4 by considering different number of days (1 week interval) between two consecutive visits with positive results. Bias and MSE are minimal if positive results observed within 35 days are considered to be of the same infection. C+ and R+ represent positive result/infection at clinical and routine visits, respectively.

### Appendix B: Data application

#### B.1 Interval-censored infection times

Interval censoring occurs if the time at risk *T_AR_* is only known to lie between two time points. In the PRISM study, the time to the second, third or the *n*-th infection is only known to lie between the point the child is tested positive and the point he/she first tested negative after recovering from the previous infection. Generally, if the real time at risk *t_AR_* for the *n*-th infection of an individual of age *a* when becoming susceptible again at calendar time *t_(n−1)_*, lies between *t_L_* and *t_U_*, then the probability density function for the time at risk is given by

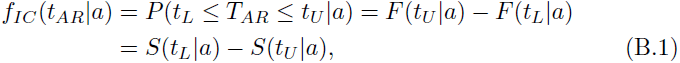

Where 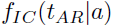is the modified density function for interval-censored data 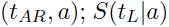and 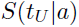 are the conditional survival functions evaluated and *t_U_*, respectively, i.e., for *t_L_*,

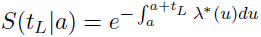

Where λ^*^(u) is the infection hazard (for symptomatic infections). In case of **exponential** infection times, we have 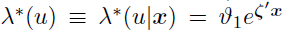and 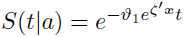, which implies

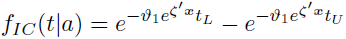

Alternatively, for the **Weibull** and **Gompertz** distributions, it is straightforward to obtain similar expressions based on the expressions for the hazard functions in Table 1 in the main text. Finally, in case of the fractional **polynomial** model, we have 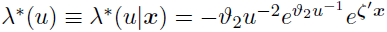and

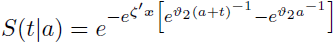

Note that in case the first event recorded for an individual of age *a* is a clinical malaria infection, the time at risk lies in the interval 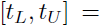
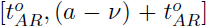 where 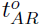 is the observed time at risk, *a* is the age of the individual at the entry of the study, and 0 ≤ *ν* ≤ *a* is the age of the individual when becoming susceptible after the last infection prior to the inclusion into the study, thereby giving rise to a contribution 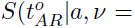
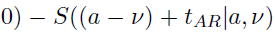 to the likelihood function. Since *ν* is unknown, we need to marginalize over the probability density function of the random *V* variable. However, this leads to complicated expressions for the likelihood function, hence, in this manuscript, we take 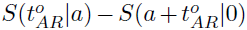 as likelihood contribution, implying that 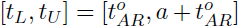, and we consider the aforementioned marginalization strategy as further research which is beyond the scope of this paper. Hereunder, we describe 4 possible situations for the treatment of interval censoring in the PRISM study. First, let *t_(*n*)_* be the calendar time at which one tests positive for the *n*-th infection (*n* > 1), *t_(*n* − 1)_* the point at which one first tests negative from the (*n* − 1)-th infection, and 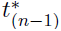 be the calendar time one was last observed positive for the (*n* - 1)-th infection.

**Situation 1:** If *t_(n−1)_* and *t_(n)_* are exactly the points when one becomes susceptible and infected, respectively, then time at risk, *t_AR_ = t_(n)_ t_(n−1)_*. In this case there is no interval censoring and the contribution to the like-lihood is simply 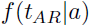, where *a* is the age of the individual at time *t_(n−1)_*.

**Situation 2:** If *t_(n−1)_* is exactly the point when one becomes susceptible, then the time at risk, 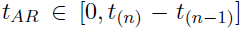, meaning that *t_L_* = 0 and 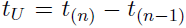 Consequently, *a* represents the age of the individual at time *t_(n−1)_* in likelihood contribution (B.1).

**Situation 3:** If *t(_n_*) is exactly the point when one becomes infected, then the time at risk, 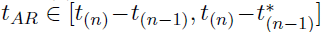, meaning that *t_L_ = t_(n)_ t(n−1*), 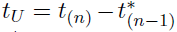 and a represents the age of the individual at calendar time 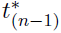

**Situation 4:** If 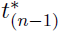 is exactly the point when one becomes susceptible, then the time at risk, 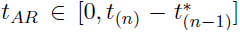, meaning that *t_L = 0_*, 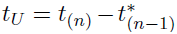and *a* represents the age of the individual at calendar time 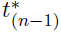

The statistical analysis presented in this paper is based on Situation 2, though the other situations are also plausible and worth considering, albeit that these scenarios are all approximations of the thruth. The impact of assumming Scenarios 3-4 on inference was found to be minor and the conclusions did not change.

#### B.2 Fit statistics

**Table B.1.** Fit statistics for models fitted to PRISM data based on Scenario 2 and 4 by study site. Better fits for each site and scenario based on AIC are indicated in bold.

| Site | Fit statistic | Exponential | Weibull | Gompertz | Fractional polynomial |
| --- | --- | --- | --- | --- | --- |
| **Scenario 2:** |  |  |  |  |  |
| Walukuba: | AIC | 1902.6 | 1867.7 | **1867.0** | 1867.8 |
|  | BIC | 1921.9 | 1889.8 | 1889.1 | 1889.8 |
| Kihikihi: | AIC | 6446.5 | 6440.1 | **6411.1** | 6492.6 |
|  | BIC | 6465.2 | 6461.5 | 6432.5 | 6513.9 |
| Nagongera: | AIC | 9864.8 | **9863.0** | 9866.0 | 9889.1 |
|  | BIC | 9883.6 | 9884.4 | 9887.4 | 9910.5 |
| **Scenario 4:** |  |  |  |  |  |
| Walukuba: | AIC | 2012.0 | 2008.3 | **1992.4** | 2092.3 |
|  | BIC | 2039.6 | 2049.0 | 2025.6 | 2122.7 |
| Kihikihi: | AIC | 6683.8 | 6028.1 | **5975.8** | 7345.5 |
|  | BIC | 6710.5 | 6060.2 | 6007.9 | 7384.1 |
| Nagongera: | AIC | 9554.4 | 9327.2 | **9304.6** | 10373 |
|  | BIC | 9581.1 | 9359.3 | 9336.6 | 10403 |

